# Analysis of Multisensory-Motor Integration in Olfactory Navigation of Silkmoth, *Bombyx mori*, using Virtual Reality System

**DOI:** 10.1101/2021.09.14.460318

**Authors:** Mayu Yamada, Hirono Ohashi, Koh Hosoda, Daisuke Kurabayashi, Shunsuke Shigaki

## Abstract

Most animals survive and thrive due to navigation behavior to reach their destinations. In order to navigate, it is important for animals to integrate information obtained from multisensory inputs and use that information to modulate their behavior. In this study, by using a virtual reality (VR) system for an insect, we investigated how an adult silkmoth integrates visual and wind direction information during female search behavior (olfactory behavior). According to the behavioral experiments using the VR system, the silkmoth had the highest navigation success rate when odor, vision, and wind information were correctly provided. However, we found that the success rate of the search significantly reduced if wind direction information was provided that was incorrect from the direction actually detected. This indicates that it is important to acquire not only odor information, but also wind direction information correctly. In other words, Behavior was modulated by the degree of co-incidence between the direction of arrival of the odor and the direction of arrival of the wind, and posture control (angular velocity control) was modulated by visual information. We mathematically modeled the modulation of behavior using multisensory information and evaluated it by simulation. As a result, the mathematical model not only succeeded in reproducing the actual female search behavior of the silkmoth, but can also improve search success relative to the conventional odor source search algorithm.

## Introduction

In many organisms, including humans, appropriate behavior is determined based on the integration of different kinds of information from the environment. Examples of information obtained from the environment include light, sound, odor, and wind. Odor, unlike sound and light, does not have high temporal or spatial resolution, but is highly persistent and diffusive, and is therefore widely used as a communication tool by organisms (***Renou, 2014***). Insects, in particular, communicate extensively using odor (e.g., aggregation pheromone, trail pheromone, sex pheromone; (***Wyatt, 2014***)), despite their small-scale neural systems. Odor information is also largely used to locate feeding sites and flowers (***Renou, 2014***).

Understanding odor-based search behavior is of great value not only in biology, but also in engineering research. This is because odor-based search can be applied to gas leak source search robots or lifesaving robots in a disaster area instead of dogs. There have been many attempts to model the search behavior of living organisms using odor and to implement (***Chen and Huang, 2019***) it in robots, but artificial systems have not yet demonstrated the same capabilities as living organisms. One of the reasons for this lack of performance is that models do not incorporate when and under what conditions, and which sensory information is fed back to inform subsequent behavior. To solve this problem, it is necessary to measure behavioral changes in insects when multiple types of sensory information are presented to them. Previously, Pansopha *et al*. (***Pansopha et al., 2014***) found that the mating behavior of a silkmoth, which is elicited by odor stimulus, was modulated by visual stimulus. Further, in a study on crickets, Haberkern *et al*. (***Haberkern and Hedwig, 2016***) reported that long-term tactile stimulation suppressed phototaxis, suggesting that behavioral switching between proximal environmental information and phototaxis may occur. These results revealed that behavioral modulation and switching mechanisms occur in all insects through the acquisition of multiple types of environmental information. However, because these experimental results were obtained when controlled stimuli were provided, the mechanisms of behavioral modulation and switching when complex environmental changes were presented are still unknown. In recent years, the use of virtual reality (VR) systems in insect behavioral experimental setups has attracted attention as a way to present complex environmental changes. For example, Kaushik *et al*. (***Kaushik et al., 2020***) found that dipterans use airflow and odor information for visual navigation using an insect VR system. Because VR systems allow for the presentation of more natural environmental changes, they also allow for quantitative analysis of the effect of behavior modulation and switching mechanisms on functions such as search and navigation. Furthermore, they allow researchers to create and test insects in situations that do not occur in nature, and may therefore play an important role in the construction of robust behavioral decision algorithms for unknown environments.

In order to elucidate the adaptive odor source search behavior of insects, we constructed a VR system that can present multiple types of environmental information simultaneously and continuously, and employed it to clarify how sensory information other than odor is used. Our VR system was connected to a virtual field built on a computer, and presented odor, wind, and visual stimuli according to environmental changes in the virtual search field. We employed an adult male silkmoth *Bombyx mori* as our measurement target. Female search behavior of the silkmoth has a stereotyped pattern (***Ryohei et al., 1992***), but the duration and speed of the behavior are regulated by the frequency of odor detection and the input of other sensory information (***Pansopha et al., 2014***; ***Shigaki et al., 2019a***,b). However, previous studies have observed silkmoth behavior in response to a specified, unchanging amount of stimulus, and the nature of behavioral changes during an actual odor source search warrants further investigation.

In this study, we used a VR system to clarify which sensory organs the silkworm moth uses to search for females, and how they are used. In addition, we constructed a model from our biological data and tested the validity of the model using a constructive approach.

## Results

In this study, we analyzed changes in female search behavior of an adult male silkmoth in response to multiple sensory inputs. To measure behavior, we constructed a novel virtual reality device that presents odor, visual, and wind stimuli (Fig. 1A) to provide the silkmoth with the illusion that it was searching for a female (see Supplemental video 1). The providing for each sensory input is defined as shown in Fig. 1B. The odor discharge port is integrated with a tethered rod for fixing the body, and the sex pheromone is discharged independently from the upper part of the left and right antennae. The direction of wind stimulation is determined by the heading angle with respect to the windward direction from the four directions: front, back, left and right. Moreover, the visual stimulus provides an optical flow in the direction opposite to the turn direction. The VR device was connected to a virtual field built on a computer, and the odor puffs in the virtual field were produced by image processing of the actual odor diffusion using an airflow visualization system (***Yanagawa et al., 2018***). In addition, the virtual field was set with wind flowing uniformly from left to right. The search performance of the silkmoth using the constructed VR system was the same as that in free-walking experiments (see Supplemental Materials). We carried out experiments using the VR by changing the number of sensory inputs as follows (Fig. 1C);

**Figure 1.**
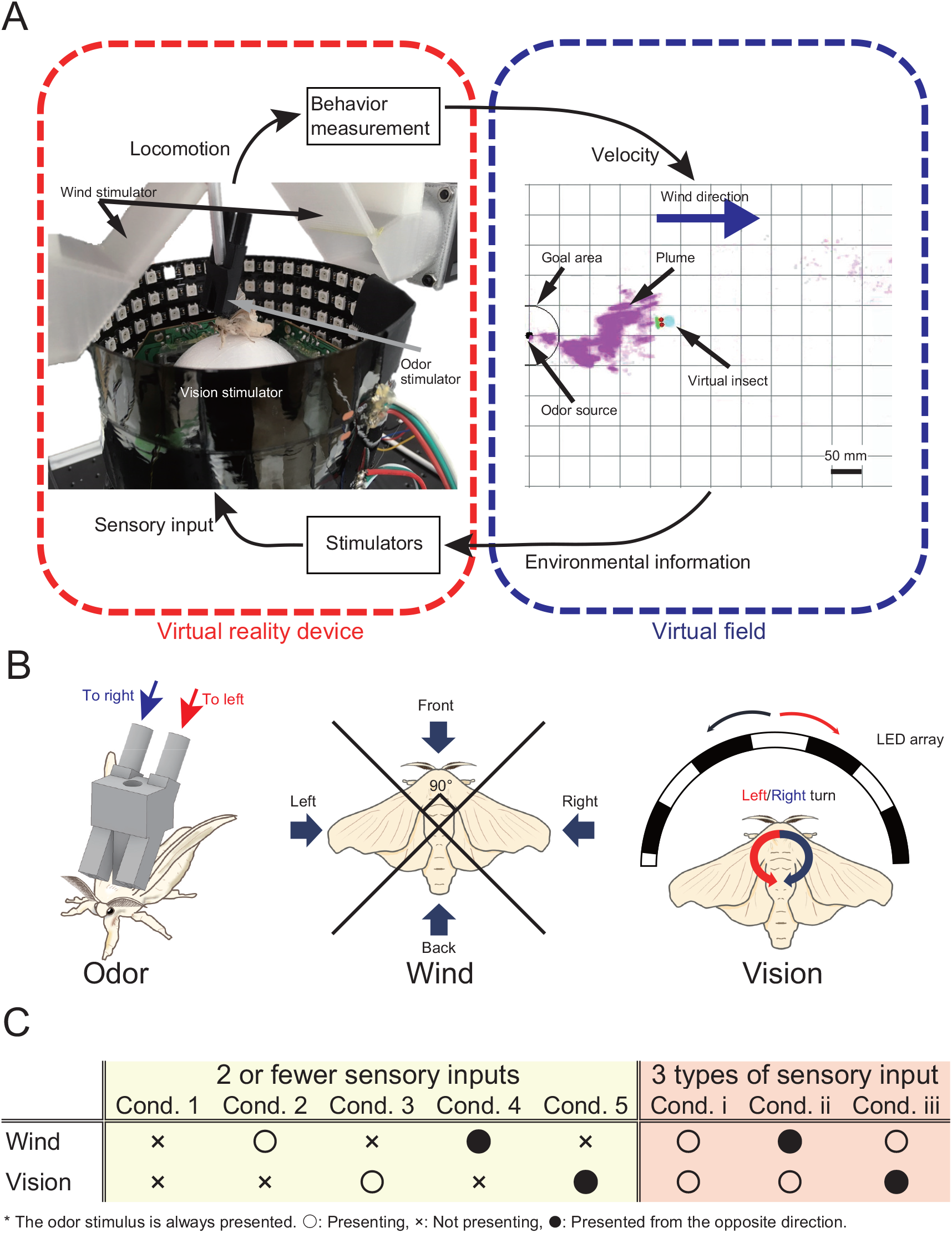
The virtual reality (VR) system for olfactory navigation of the insect and a list of experimental conditions. A: The VR system is equipped with a stimulator of odor, vision, and wind, and is connected to a virtual odor field. The insect on the VR device performs olfactory navigation in virtual space. B: Definition of presentation way of each sensory stimulus. C: The odor is presented under all conditions. “⚬”, “×”, and “•” indicate presented, not presented, and presented from the opposite direction to the actual direction, respectively. **Figure 1–Figure supplement 1**. System configuration diagram and evaluation of VR system. **Figure 1–video 1**. A video of a silkmoth behavior experiment using a VR system.

- Presents two or less types of sensory input (Group1: odor only / odor + wind / odor + vision),
- Presents three types of sensory input (Group2: odor + wind + vision).

“⚬” indicates that we presented the environmental information to the silkmoth, and “×” indicates that we did not present the environmental information to the silkmoth (Fig. 1C). “•” indicates that the stimulus was presented from the opposite direction. For example, “Inverse wind” means that the virtual insect in the computer received wind from the front, while the real silkmoth received wind from the back. Three repetitions of the experiment were conducted using 10 silkworm moths for each environmental condition (*n*=30). We set a time limit of 300 seconds because theoretically, the silkmoth could reach the odor source in an infinite time. The search was considered a failure if the moth did not enter a radius of 10 mm from the odor source within the time limit.

### Wind and visual effects on the olfactory navigation

We compared the search performance in response to different types of stimuli by presenting wind and visual stimuli in addition to olfactory stimulus in all conditions. We first measured the behavior when wind or visual stimuli is providing in addition to odor stimulus (Group 1). We created a migration probability map in order to visualize the effects of differences in environmental conditions on the behavioral trajectory (Fig. 2A—E). To quantitatively evaluate the similarity between the migration probability maps, we calculated the earth mover’s distance (EMD) (***Rubner et al., 1997***). EMD is an index that calculates whether the distributions of two histograms are similar. The smaller the EMD value, the more similar the two histograms, and the larger the value, the less similar. Fig. 3F shows the result of calculating EMD based on cond. 1, in which only the odor stimulus was presented. The silkmoths’ trajectory in cond. 3 and 5 (odor + vision) was very similar to the trajectory in cond. 1 (only odor), suggesting that vision did not significantly affect search behavior. In addition, the EMD value when wind stimulus was presented in addition to odor was *>* 10, suggesting that the addition of wind stimulus did significantly affect search behavior. We illustrated the navigation success rate and search time (yellow background in Fig. 3A and B). Additionally, we calculated the search success rate per unit time (SPT) as a measure of the relationship between the search success rate and search time (yellow background in Fig. 3C). Higher SPT values represent better search performance. It was found that the success rate was most improved when the wind was presented correctly (cond. 2) compared to when only the odor stimulus was provided (cond. 1). The success rate was lower under the condition where the wind was presented from the direction opposite to the direction detected in the virtual environment (cond. 4), compared with the condition where only odor stimulus was presented (cond. 1). Moreover, SPT in cond. 3 and 5, in which the direction of the visual stimulus was changed without presenting the wind stimulus, were almost the same (0.411 vs. 0.409), suggesting that visual stimulus had little effect on the search performance. From this, it was found that the success rate of navigation tends to improve by correctly presenting wind stimulus in addition to the odor.

**Figure 2.**
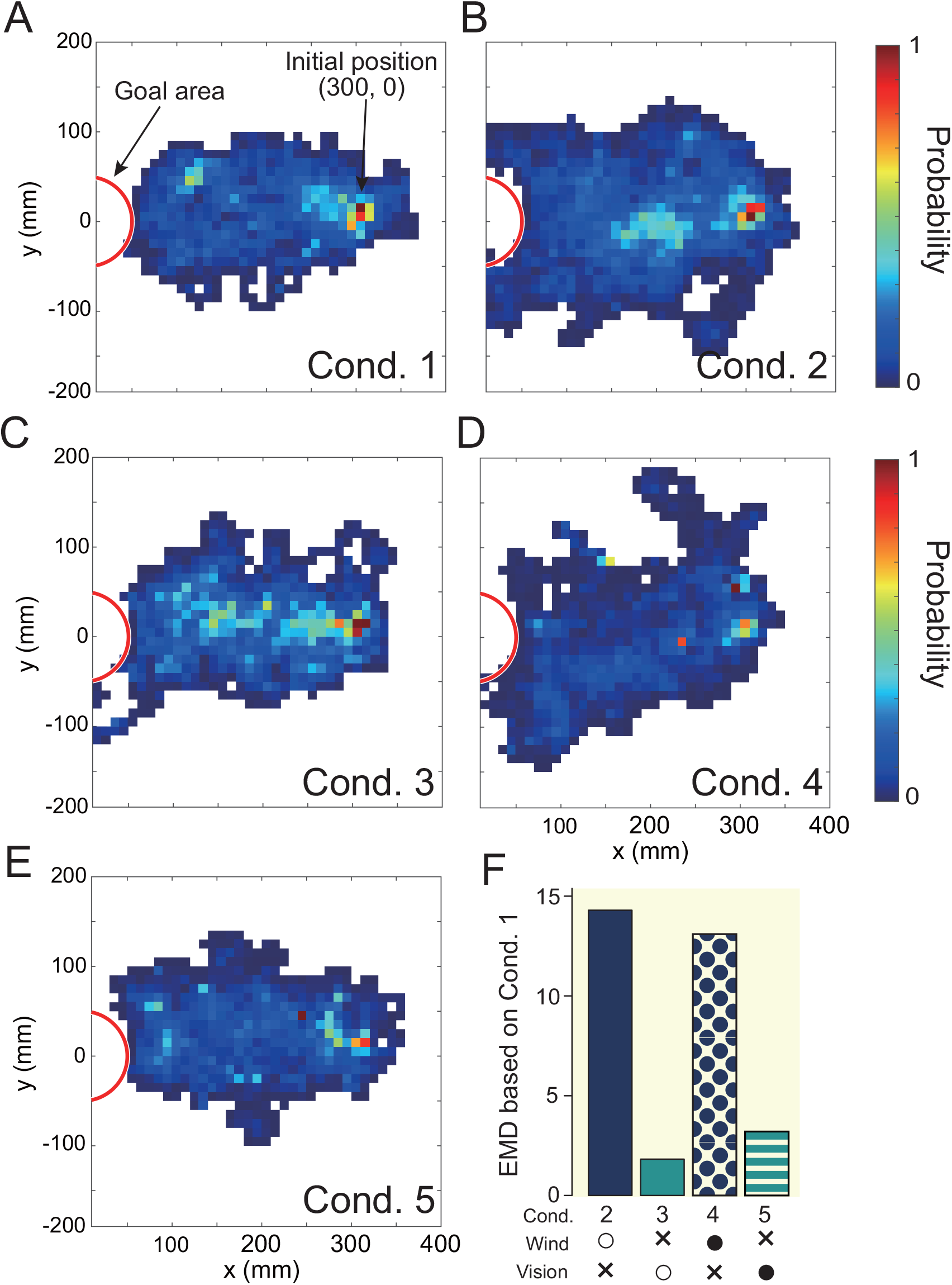
The result of visually expressing the trajectory under each experimental condition with a migration probability map (A—E). The white area in the figure indicates that the moth did not move in that space. A quantitative evaluation using earth mover’s distance, EMD is illustrated in F. F was the result of calculating the similarity (EMD) based on the trajectory of cond. 1 (odor presentation only). The lower the value, the higher the similarity, and the higher the value, the lower the similarity of the trajectories.

**Figure 3.**
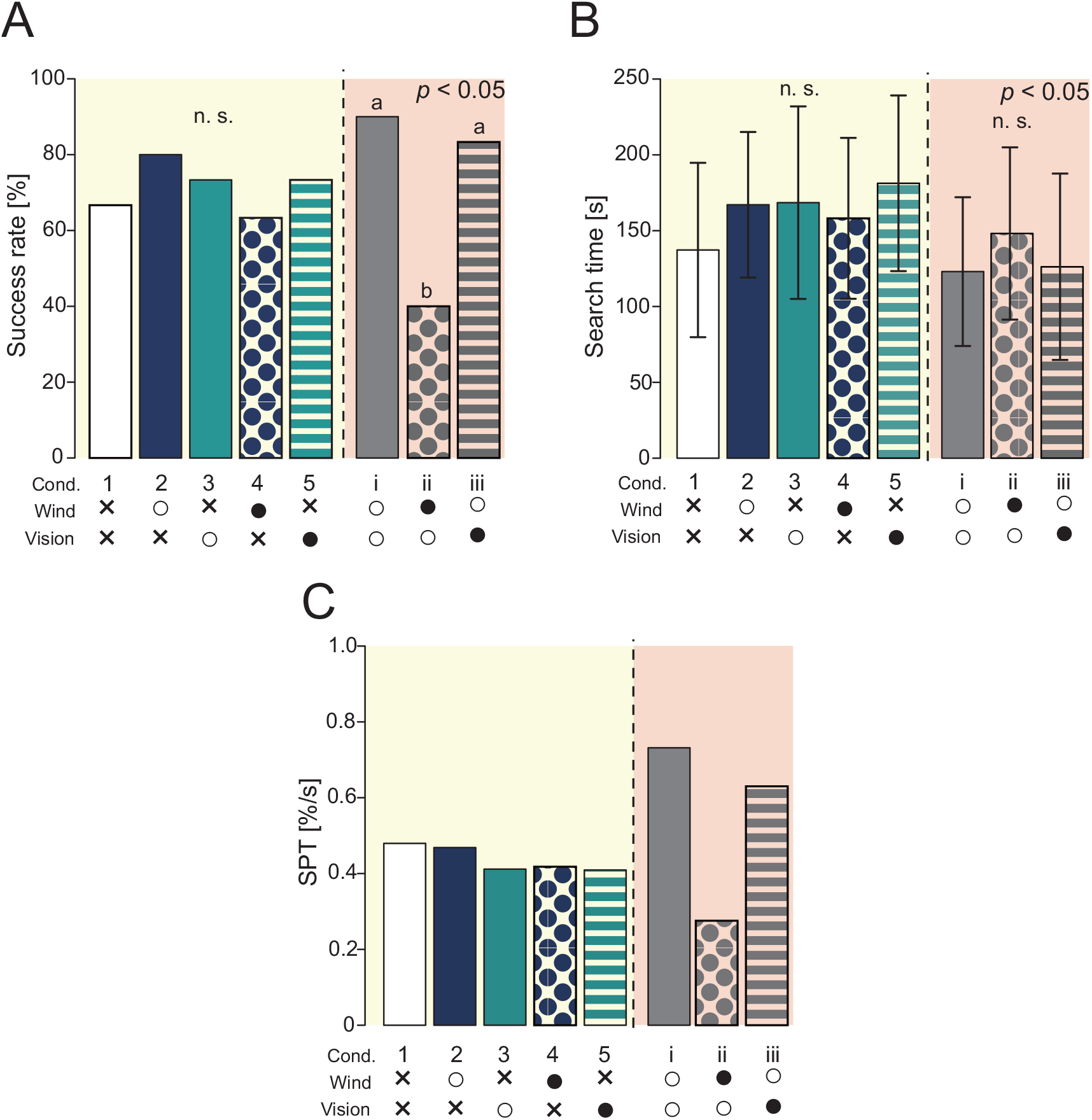
Results of navigation experiments using virtual reality (VR). The yellow background is the result of providing 2 or fewer sensory stimuli, and the red background is the result of providing 3 sensory stimuli. A: The success rate of the navigation (Fisher’s exact test, *p <* 0.05). B: The search time at the time of success (Steel-Dwass test, *p <* 0.05). C: Search performance. The success rate per unit time was calculated based on the results of A and B.

We found that when three types of sensory stimuli were presented (Group2), the success rate changed significantly compared to when two types of sensory stimuli were presented (red background in Fig. 3). Focusing on cond. i, the SPT values were also significantly different, suggesting that the olfactory navigation can be performed more accurately and effciently by presenting all sensory stimuli. Additionally, under the condition that the wind stimulus is presented from the direction opposite to the direction received in the virtual environment, the search success rate is significantly reduced, and it is clear that the wind direction information is an important factor. Together, we hypothesized that the silkmoth has achieved effcient female search using all sensory inputs: odor, wind, and vision.

### Extraction of behavioral modulation mechanisms in the odor source search

In the previous section, it was found that wind direction information, in addition to odor information, contributed to improving the success rate in searching for the odor source. Here, we analyze in detail how behavior is modulated by visual and wind information using three experimental conditions: a forward condition (cond. i), an inverse condition for wind direction information (cond. ii), and an inverse condition for visual stimuli (cond. iii).

We first analyzed the effect of wind direction information on odor source search behavior. The results of calculating EMD in the previous section suggested that the search behavior was significantly affected when wind direction information was presented, in addition to odor information. Because the key stimulus for female searching behavior in silkmoths is odor, and because previous studies (e.g. (***Kikas et al., 2001***)) have reported that odor detection frequency is related to the distance from the odor source, and that odor detection frequency modulates behavior, we analyzed odor as a state variable. Here, odor detection frequency is defined as the frequency at which the silkmoth receives odors per unit of time. Fig. 4A shows a boxplot of the relationship between odor detection frequency and movement speed when wind direction information is presented in the forward direction. Fig. 4B shows the results when the wind direction information in the virtual and real environments is presented in opposite directions. The black dot in the white box represents the average value. In Fig. 4A and B, the horizontal axis represents odor detection frequency, and the vertical axis represents the translational and angular velocities. The red box plot in the figure shows the peak velocity. When the direction of the wind in the virtual environment and the direction of the wind presented to the silkmoth in the real environment matched, odor detection frequency peaked at 0.7 Hz. However, in the non-matching condition, the silkmoths always moved at a constant speed, regardless of the odor detection frequency. The angular velocity peaked at 0.4 Hz in the matched condition but shifted to 0.2 Hz in the non-matched condition. The peak translational velocity was very close to the frequency of female sex pheromone emission (approximately 0.8 Hz (***Fujiwara et al., 2014***)). When odor detection frequency was low, the female was far from the male, and therefore the male searched actively. However, as the distance from the female decreased, the frequency approached 0.7 Hz, which suggests that the male may be trying to locate the female more carefully by increasing the spatiotemporal resolution of the search. On the other hand, when the wind and odor directions do not coincide, the silkmoth may try to stay within the reach of the odor by constantly searching regardless of the odor detection frequency. This may be because the peak of the angular velocity shifts toward a lower odor detection frequency, and if the odor and wind direction do not match, the silkmoth rotates slowly to carefully search for the odor.

**Figure 4.**
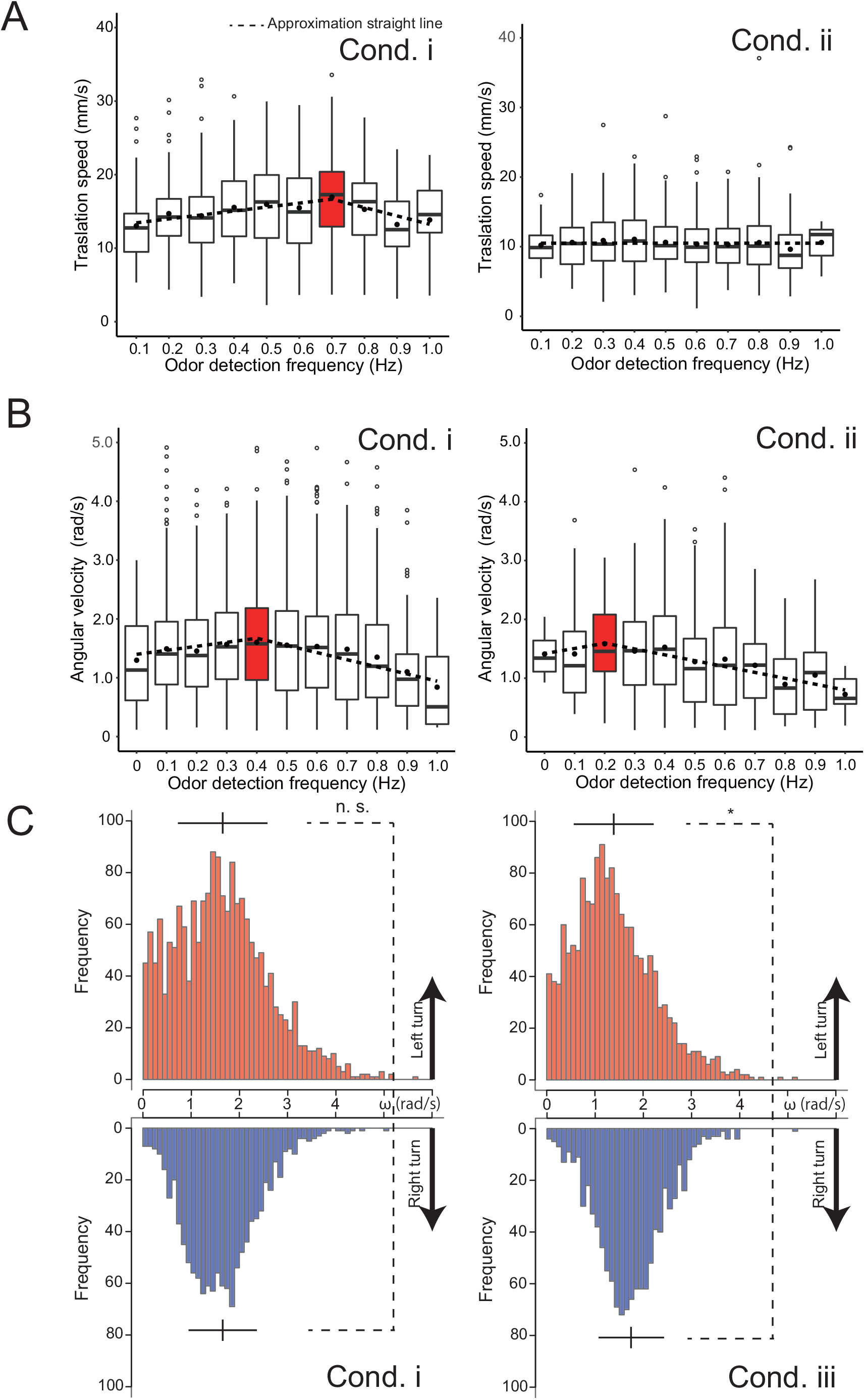
Analysis of the effects of wind and vision direction on behavior. A: Changes in the translational velocity and angular velocity when the wind direction was presented correctly (cond. i). Both translation and angular velocity peaked, and the behavior was modulated depending on the odor detection frequency. B: Changes in translational velocity and angular velocity when the wind was presented from the direction opposite to the actual direction (cond. ii). The translational velocity was constant regardless of the odor detection frequency, and the peak position of the angular velocity was 0.5 times that of cond. i. C: Comparison of angular velocities when the visual information was presented correctly (cond. i) and when it was presented in the opposite direction (cond. iii). Vision was used to equalize the speed of the left-right rotation.

Next, we investigated the effects of visual stimuli on odor source search behavior. Section 2.1 suggests that visual stimuli do not directly contribute to search performance. However, many flying insects with compound eyes use visual information for their own postural control (***Dyhr and Higgins, 2010***)(***Dyhr et al., 2013***). The silkmoth, although a walking insect, may retain some vestiges of flying insects (***Kanzaki, 1998***)(***Shigaki et al., 2016***). Therefore, we hypothesized that vision may be used for postural control. To test this hypothesis, we compared the angular velocities of silkmoths in cond. i (all stimuli in the forward direction) to those in cond. iii (visual stimuli presented in the inverse direction). Fig. 4C shows a histogram of the left and right angular velocities under each experimental condition. The red color in Fig. 4C indicates the angular velocity when rotating counterclockwise, and the blue color shows the angular velocity when rotating clockwise. The position of the vertical bar above the histogram represents the average angular velocity, and the length of the horizontal bar represents the standard deviation. Although the average angular velocities of the left and right rotations were the same in cond. i (visual stimuli presented correctly), the angular velocities of the left and right rotations differed when the visual stimuli were presented in the opposite direction to reality (Wilcoxon rank sum statistical test, *p <* 0.05). These results indicate that silkmoths, like flying insects, use visual stimuli for postural control.

We found that the silkmoth modulated its behavior based on whether or not the direction of odor and wind detection coincided.

### Modeling and validation of behavioral modulation mechanisms

Here, we investigated how behavioral modulation obtained from behavioral experiments using a VR system contributes to the odor source search. A silkmoth moves by walking on a two-dimensional plane with six legs, but it does not move in the lateral direction. Therefore, we assumed that it has non-holonomic constraints and constructed our model to output straight-ahead and rotational movements. Previous studies have proposed a silkmoth search model called the surge-zigzagging algorithm (***Ryohei et al., 1992***)(***Shigaki et al., 2019b***), which makes action decisions using only olfactory information. In this study, we updated the surge-zigzagging algorithm to incorporate wind direction information. For convenience, we called this algorithm MiM2 the multisensory input-based motor modulation) algorithm. A block diagram of the constructed MiM2 algorithm is presented in Fig. 5A. The model has a straight-ahead speed (*v*) controller (1) and an angular velocity (*ω*) controller (2), and each controller changes output depending on the amount of sensory information input. The detailed equations for each controller are shown below:

**Figure 5.**
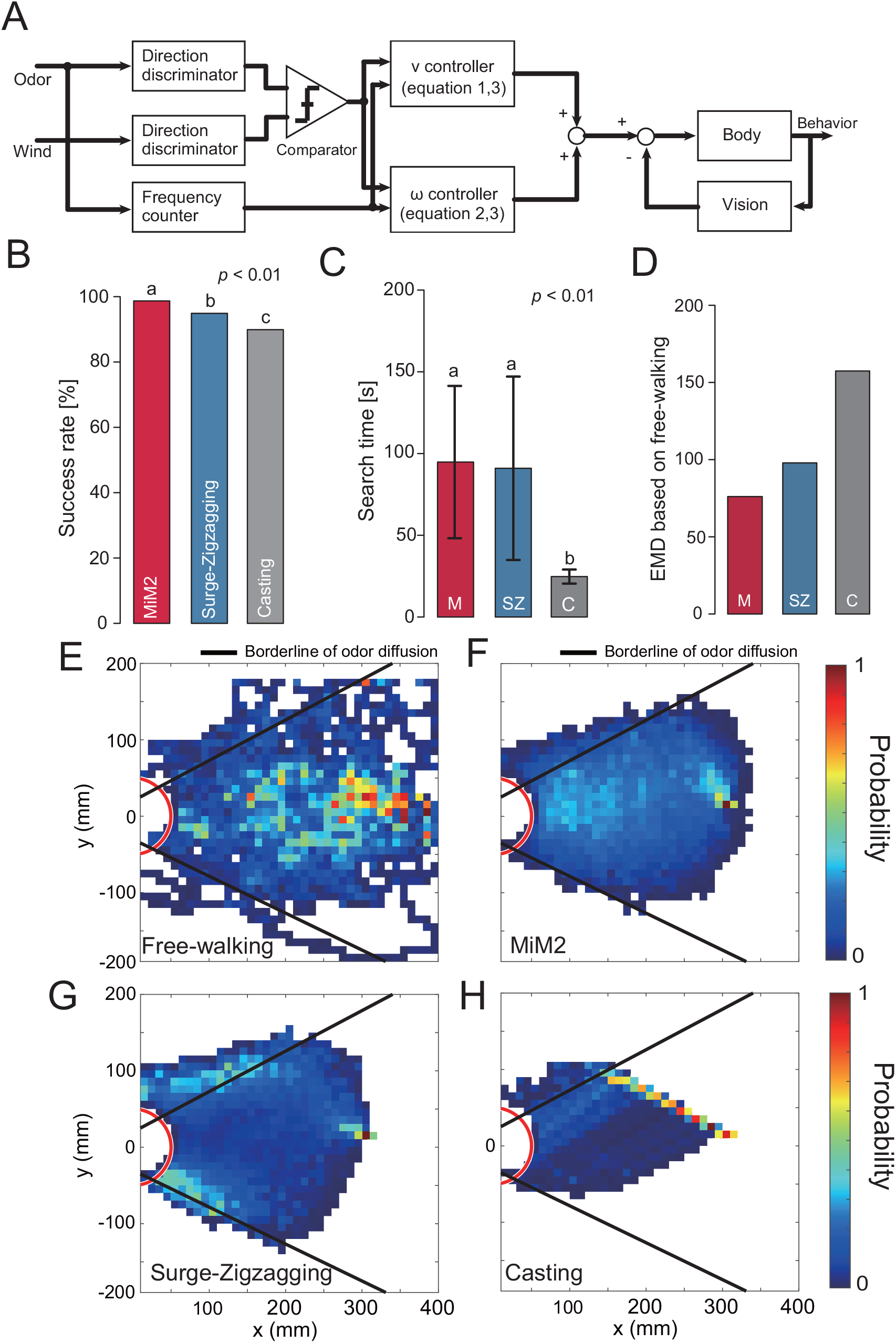
Block diagram of the proposed model and the simulation results. A: Cont2rol model that modulates speed according to the degree of coincidence between odor detection direction and wind direction. B: The success rate of navigation. C: The search time at the time of success. D: The EMD of each search algorithm calculated based on the migration probability map from the free-walking experiment. E—H: The migration probability map at the time of success. **Figure 5–Figure supplement 1**. Flowchart of other algorithms. **Figure 5–video 1**. Experimental video of the simulation.

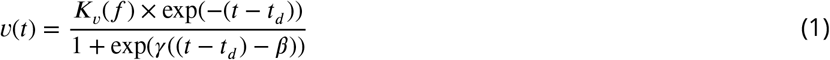

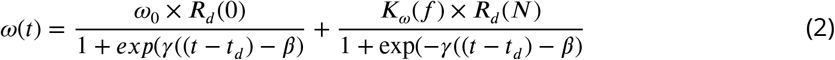

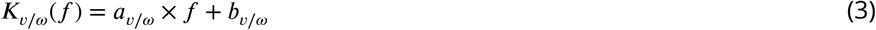

The free parameters were set to *γ* = 1000, *β* = 0.50, and *ω*_0_ = 0.57. Here, *f* denotes the odor detection frequency. Moreover, *K*(*f*) is a linear function that varies with the direction of odor detection and wind detection. The parameters a and b of *K*(*f*) are shown in Table 1. These were obtained by approximating the results of the behavioral experiment to the least-squares method (Dotted line in Fig. 4A, B). In addition, *t*_*d*_, *R*_*d*_, and *N* represent the odor detection timing, the odor detection direction, and the number of turn motions, respectively. *R*_*d*_ (0) represents a straight-ahead motion state and takes a value of -1 when the left antenna detects and 1 when the right antenna detects. If both antennae detect, a value of -1 or 1 is randomly selected. *N* increases according to Equation (4) up to four, but does not take a value of four or more because the number of zigzagging motions was approximately three in a previous study (***Ryohei et al., 1992***). When it receives an odor stimulus according to Equation (1–3), it generates a straight motion for 0.5 seconds and then makes a turn motion.

**Table 1.**
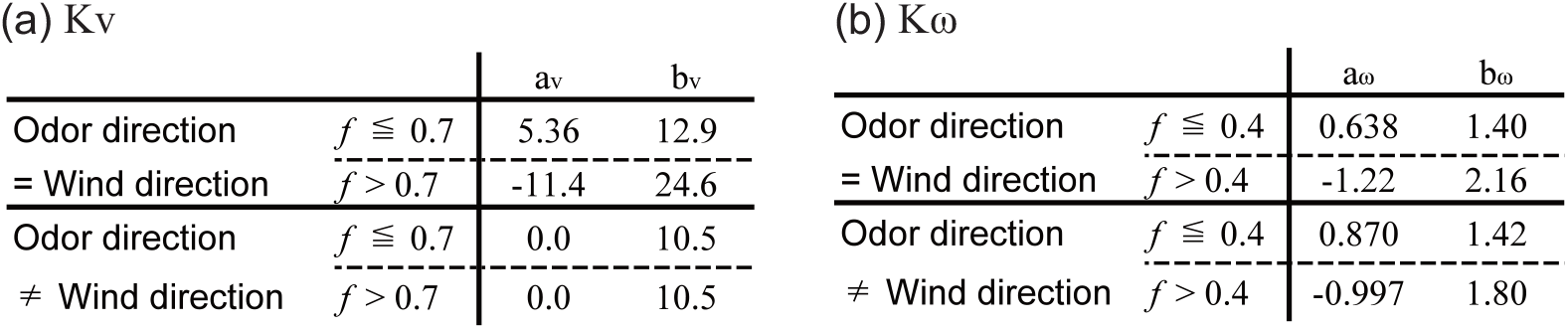
List of parameters for *K*(*f*).

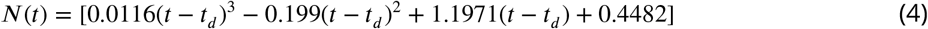

By passing through the directional discriminator and frequency counter, the odor is converted into information such as whether it was detected by the left or right antennae and how much odor it was exposed to. The wind is converted into wind direction information. A comparator is used to determine whether the direction of odor detection and the direction of the wind are the same, and the results are input to the straight-ahead speed and angular velocity controllers, which also receive odor detection frequency. Visual information controls the angular velocity of the left and right rotational movements following which olfactory behavior takes place. From this information, the movement speed output is calculated, and the behavior is generated. We compared the MiM2 algorithm to the previous surge-zigzagging algorithm and the casting algorithm, which uses wind information for searching in a moth-inspired algorithm proposed in a previous study (***Li et al., 2001***).

The simulation environment employed a virtual environment similar to that used in the behavioral experiments in the VR system (see Supplemental video 2). For each algorithm, we performed 1000 odor source search experiments to evaluate search success rate and trajectory. The results are shown in Fig. 5B, C and D. Fig. 5B and C show the search success rate and the search time, respectively. MiM2 algorithm showed a significantly higher search success rate than the other algorithms (Fisher’s exact test, *p <* 0.05). The casting algorithm showed the shortest search time (Steel-Dwass test, *p <* 0.05) because it is an algorithm that actively moves upwind using odor detection. However, if the heading angle when moving upwind was incorrect, the search success rate of this algorithm was the lowest of the three, because there was a high possibility that the agent would have moved in the wrong direction. Next, we focus on the trajectory shown in Fig. 5E—H. The black line in Fig. 5E—H shows the range that the odor has a high probability of reaching. The search trajectory shows that the MiM2 algorithm always placed the searching agent in the middle of the odor distribution, whereas the conventional surge-zigzagging and surge-casting algorithms tended to place the searching agents near the edges of the odor. We evaluated the differences in trajectory between algorithms quantitatively by calculating their EDMs (Fig. 5D). The comparison of these EDM values suggested that the MiM2 algorithm most closely resembled actual silkmoth movements, and was the closest to the results of behavioral experiments under the free walking condition. Thus, the MiM2 algorithm can simulate search behavior similar to that of real silkmoths by modulating the movement speed based on wind information in addition to odor and equalizing the left-right angular velocities based on visual information.

## Discussion

### Virtual reality for insect behavior measurements

All animals, including insects, need to navigate for survival and reproduction. It is especially important to navigate effciently in harsh environments and in situations in which there are many competitors. Previous studies have investigated the homing behavior of a desert ant (*Cataglyphis*) (***Wehner, 2003***), the pheromone source localization behavior of a male silkmoth (*Bombyx mori*) (***OBARA, 1979***), and the sound source localization behavior of female crickets (*Gryllus campestris L*.) (***Schmitz et al., 1982***). These studies have contributed to our understanding of the sensory-motor integration mechanism that converts sensory input into motor output, an important function of the neural system. However, the insects in these studies may not have displayed navigation behavior as they would have under natural conditions, because stimuli were controlled and presented in a fixed amount and at regular intervals in the behavioral experiments. In addition, the nature of behavior change in a large-scale space and over a long period is relatively unknown because the behavioral experiments were carried out in a relatively small space and over a relatively short time. Because of this, multimodal virtual reality (VR) systems have attracted a great deal of attention (***Kaushik et al., 2020***)(***Naik et al., 2020***). The advantage of using a multimodal VR system is that not only can behavior be measured at a large spatiotemporal scale, but the transmission of stimuli can be precisely controlled and the behavioral output can be precisely measured. Biological data describing the relationship between sensory input and motor output are useful not only for clarifying biological functions, but also for the field of robotics, because these data play a very important role in modeling insect systems. Because the multimodal VR system in the previous studies was developed for visual-dominated navigation, it was difficult to measure olfactory-dominated navigation, which was the focus of our research. Therefore, we developed a novel multimodal VR system that allowed us to measure changes in navigation behavior when the olfactory, visual, and wind directions of the silkmoth were modified. Conventionally, previous studies on the silkmoth have shown that (1) a “mating dance” was elicited in response to sexual pheromones (***OBARA, 1979***), (2) the condition in which visual stimuli was presented immediately after the reception of sex pheromones influenced the subsequent rotational behavior (***Pansopha et al., 2014***), and (3) behavioral inhibition occurred in response to frontal winds (***Shigaki et al., 2019a***). However, the integration of the above phenomena during navigation has not been investigated. By presenting three types of sensory information simultaneously as well as continuously, we were able to clarify the roles of each type of sensory information in navigation. Moreover, mathematical modeling and simulation showed that multimodal sensory information improved silkmoth navigation.

### Behavioral modulation to odor frequency

Organisms of all size scales rely on the ability to locate an odor source in space. The chemotaxis of bacteria and a nematode have been determined by measuring and analyzing the relationship between chemical stimulus input and behavioral output under a controlled environment (***Berg, 2008***)(***Lockery, 2011***). This study found that as the concentration of chemical stimuli increased, the probability of rotating behavior decreased linearly. In other words, bacteria or nematode chemotaxis followed the odor gradient, which allowed them to locate the odor source. In the space where bacteria and the nematode exist, chemotaxis is effective in part because the odor field of the environment does not change significantly due to wind. In the case of larger-scale animals, the odor field of the environment is quite complex and it is difficult to reach the odor source by following the gradient alone. Odor fields can be complex because odor molecules are transported by airflow and mixed with other molecules at their destinations, forming complex structures (***Crimaldi and Koseff, 2001***)(***Murlis et al., 1992***). In addition, the odor molecules themselves are discrete in space, and do not carry information about the source of the odor. However, a study using a sensor array to measure the arrival of odors carried by the wind revealed that the odors emitted from the source arrive periodically (***Kikas et al., 2001***)(***Murlis et al., 2000***). Moreover, the periodicity is correlated to some extent with the distance from the source; the closer the source, the shorter the cycle, and the farther the source, the longer the cycle of arrival. Neither the concentration of the odor nor the number of exposures played an important role in the search process in *drosophila*, but the “tempo” at which flies encountered the odor was an important factor in the decision-making process (***Celani, 2020***)(***Demir et al., 2020***). If we assume that the “tempo” is the rate of odor detection per unit time, it is related to the cycle in which the odor arrives. We hypothesized that the silkmoth, like flying insects such as *drosophila*, modulates its behavior based on the “tempo” of the odor, therefore we included odor frequency in our analyses.

When the direction of the wind and the odor coincided, movement speed peaked when the odor frequency was 0.7—0.8, which is similar to the frequency at which a female silkmoth releases sex pheromones (0.79 ± 0.05 Hz) (***Fujiwara et al., 2014***). Based on these findings, we can hypothesize that if the male silkmoth detects wind and odor from the same direction, it correctly moves in the direction of the female and actively searches the field until it reaches a location where the sex pheromone release frequency approximates that of the female. When the frequency exceeded 0.8 Hz, the male seemed to be in the vicinity of the female and therefore increased the spatiotemporal resolution of the search in order to locate the female and prepare for the transition to mating behavior. This might correspond to the silkmoth switching to the odor source declaration algorithm in olfactory navigation. The biological experimental data obtained in this study are consistent with those of earlier research demonstrating that an algorithm that shortens the travel distance of the surge toward the end of the search improves the search performance of a robot (***Shigaki et al., 2017***).

In an earlier study investigating the direction of wind and odor in the environment, the direction of wind and odor were the same in the open field experiment (no obstacles) (***Murlis et al., 2000***). However, the direction of wind and odor is not always the same in a complex environment such as a forest with many trees (***Murlis et al., 2000***). Our data showed that in situations where the wind and odor direction do not match, the silkmoth always moves at the same rate given any odor detection frequency. This may be a chemical tracking strategy to avoid leaving the odor range by moving at a speed lower than normal speed, and suggests that the silkmoth is estimating the degree of environmental turbulence and modulating its behavior based on the odor and wind direction information. This strategy may be applicable to an engineering search system that switches the search strategy according to the environmental conditions.

### Comparison with conventional CPT algorithm

Odor sensing plays an important role in situations in which visual search is difficult (a space filled with darkness and thick smoke). However, because the technology for development of an odor sensor is lagging behind that of other sensory sensors (e.g., cameras, microphones), robot olfactory research is still a developing field. Currently, dogs play the role of odor sensors, but due to the high cost of training dogs and the deterioration of their odor sense with physical condition and age, engineering solutions using autonomous mobile robots for searching are needed. Conventional robot olfactory research has emphasized the development of motion planning (search algorithm). The search algorithm is roughly divided into two fields: one is a bio-inspired algorithm that imitates the search behavior of living organisms, and the other is a search that estimates the position of the odor source using a statistical method (statistical algorithm). The bio-inspired algorithm, which mimics the behavior of organisms that can search in real time, is superior to the statistical algorithm in terms of robot implementation (e.g., (***Russell et al., 2003***)(***Lochmatter et al., 2008***)). However, behavior patterns and movement speeds in these bio-inspired algorithms are always constant, regardless of the environmental conditions. Accordingly, there was a problem that the original search performance of living things could not be replicated. For this reason, different kinds of research have been carried out more recently and applied to the bio-inspired algorithm, such as information-theoretic analysis of the trajectory of an insect (***Hernandez-Reyes et al., 2021***), extraction of adaptability from neuroethology data by fuzzy inference (***Shigaki et al., 2019b***), and acquisition of behavioral switching indices in response to environmental changes by measuring insect behavior while visualizing airflow (***Demir et al., 2020***). In this study, we found that behavioral modulation occurred based on the relationship between the direction of the odor arrival and the wind, and we used our data to reproduce a behavioral trajectory that was not only better than that of the conventional bio-inspired algorithm, but was also more similar to the search behavior of the actual silkmoth. Therefore, it is clear that a probabilistic and time-varying behavior modulation mechanism has a better search performance than a time-invariant search algorithm.

## Materials and methods

### Animals

Silkmoths, *Bombyx mori* (Lepidoptera: Bombycidae) were purchased from Ehine Sansyu Co., Japan. Adult male moths were cooled at 16°C one day after eclosion to reduce their activity, and were tested within 2–7 days after eclosion. Before the experiments, the moths were kept at room temperature (25—28 °C) for at least 10 min.

### Virtual odor field

To generate a virtual odor field closely mimicking reality, smoke was emitted into the actual environment. The diffusion of the smoke was recorded and implemented into the simulator using image processing. The smoke visualization experiment was conducted in an approximately 2.5 × 0.8 m area in a darkroom. First, the smoke was visualized using a smoke generator and a laser sheet. The smoke emitted into the room gleamed when it hit the irradiated laser sheet, and the distribution of smoke was recorded using a high-sensitivity camera. In this experiment, we carried out a visualization experiment in a darkroom because it is important to accurately record the light reflected by the smoke. This method was based on particle image velocimetry (PIV), which measures the velocity of fluid flow in space by visualizing particles and analyzing their movements. PIV accuracy is improved by scattering a large amount of smoke (particles) in space because it focuses on accurately tracking each particle, however, our method did not require following each particle. Instead, our method used a mass of smoke as an odor plume, and we observed how smoke floated and was distributed in space. In the visualization experiment, the flow meter (1.0 L/min) and solenoid valve (1 Hz) were controlled under the same released conditions as in the behavioral experiment. Our method required less smoke than does PIV, and the smoke is thinner and more diffcult to visualize. For this reason, it was necessary to increase the sensitivity of the camera, however, noise increased accordingly. Moreover, in PIV, it is not necessary to distinguish between smoke particles and dust because only the movement of particles in space is important, but our method required removing other particles in order to measure the smoke position alone as the plume. Hence, we applied image processing to the smoke image. Luminance values can have a large range because the intensity of smoke varies with airflow and time. In addition, it is diffcult to extract only smoke with simple thresholding because the noise caused by dust has the same luminance as does smoke. Therefore, we adopted a method that focuses on connected components to remove noise while maintaining the shape of the smoke. In grouping based on connected components, two objects are considered connected if adjacent pixels in a binary image take the value of 1. Because smoke exists as a mass, it can be inferred that it has an area above a certain level when divided into connected components. For this reason, we removed the pixels whose connected components were not in adjacent pixels and whose area did not exceed a certain level. These processes were performed with source code using OpenCV.

### Configuration of virtual reality system

The behavioral experiment was carried out using a homemade virtual reality system (a photograph of the actual device is shown in Fig. 1A). The traditional tethered measurement system is used for behavior measurement, and a stimulator that presents odor, visual, and wind direction information is installed around the tethered measurement system (Figure 1-Figure supplement 1A). The details of each stimulator are as follows.

#### Odor stimulator

We provided sex pheromone stimulation to both antennae of a silkmoth using Bombykol ((E,Z)-10,12-hexadecadien-1-ol). A cartridge containing 1000 ng of bombykol was placed in a tube in order to present air containing bombykol to the silkmoth. The compressed air from the air compressor (NIP30L, Nihon Denko, Aichi, Japan) passed through three gas washing bottles containing degreased cotton, activated carbon, and water, respectively, and was adjusted to 1.0 L/min using a flow meter (KZ-7002-05A, AS ONE CORPORATION, Osaka, Japan). The timing of the stimulation was controlled by switching the flow path of the solenoid valve (VT307, SMC Corporation, Tokyo, Japan). The odor stimulus discharge ports were integrated with the tethered rod, and the discharge ports were located above the antennae. The tethered rod with the discharge ports was fabricated using a 3D printer (Guider2, Flashforge 3D Technology Co., Ltd., Zhejiang, China). As a result of presenting a one-shot pheromone stimulus to the silkmoth using this odor stimulator, the odor stimulus can be presented correctly because the female search behavior was elicited (Figure 1-Figure supplement 1B).

#### Vision stimulator

Because the optical flow presentation method using LED arrays was also used in previous studies with flying insects, we also use the same method in this study. The visual stimulus device was constructed by arranging LEDs (WS2812B, WORLDSEMI CO., LIMITED, GuangDong, China) with a built-in microcomputer in an array around the tethered measurement system. The LED array was comprised of 256 LEDs arranged in 8 (vertical) × 32 (horizontal) panels. The array was presented as an optical flow by controlling the lighting timing of each LED according to the angular velocity of the silkmoth.

Because it has been reported that the silkmoth causes optomotor reflexs with respect to optical flow and the neck tilts in the direction of optical flow (***Minegishi et al., 2012***), whether or not this vision stimulator is functioning was evaluated by the angle of the neck with respect to optical flow. The vision stimulator is functioning properly because the silkmoth has the largest neck tilt in the range of angular velocity during the female search behavior (Figure 1-Figure supplement 1C), and this result is similar to the past research data (***Pansopha et al., 2014***).

#### Wind stimulator

The wind presented to the silkmoths was generated using the push-pull rectifier (***González et al., 2008***). The push-pull rectifier consists of a push side that sends out the wind and a pull side that draws in the sent wind. Fans (PMD1204PQB1, SUNON, Takao, Taiwan) were installed on both the push and pull sides to generate airflow. Space can be used effectively using this device because air can flow without covering the workspace with a partition. The push-pull rectifier was connected to a hollow motor (DGM85R-AZAK, ORIENTAL MOTOR Co., Ltd., Tokyo, Japan) which rotated to generate wind. The performance of the push-pull rectifier was evaluated using particle image velocimetry (PIV) (***Adrian, 2005***). For PIV analysis, we used videos that were shot at 800 fps with a resolution of 640 × 480 pixels. As a result, it was confirmed that the wind generated by the push-pull rectifier did not generate vortices (Figure 1-Figure supplement 1D).

#### VR system evaluation experiment

We experimentally verified the extent to which a VR system equipped with odor, vision, and wind direction stimulators could reproduce a free-walking experiment in a real environment. In the free-walking experiment in a real environment, the search field was the same size as the VR system, and an odor emission frequency (1 Hz) was used. Two repeated experiments were carried out using 15 moths, and their behavior was recorded using a 30 fps video camera (BSW200MBK, BUFFALO, Aichi, Japan). The figure shows the results of the quantitative comparison of search success rate and relative distance (Figure 1-Figure supplement 1E, F). We found that there was no difference between the results of VR and the free walking experiment. In other words, VR was able to reproduce the experiment of free walking, and the behavior experiment may be performed using the VR device proposed in this study.

### Migration pathway ratio map

The migration pathway ratio map statistically processed the trajectories of all trials and denoted the points where the quadcopter frequently passed through the experimental field. We created the migration pathway ratio map *C*_*n*_(*x, y*) based on the rule given by Equation (5). Here, *n* represents the trial number, and the size of a grid cell is 0.1 × 0.1.

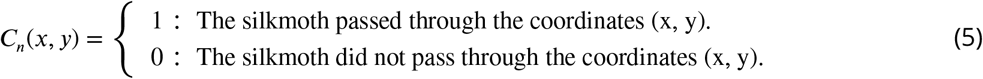

We applied the equation to all trial trajectories and summarized them, and then calculated the trial average based on equation (6) to create the migration pathway ratio map *P* (*x, y*). The migration pathway ratio map was created using MATLAB (2020a, MathWorks, MA, USA).

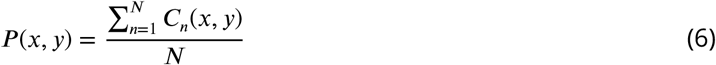

### Statistical Analysis

For all data analyses, R version 4.0.3 was used (R Core Team). All earth mover’s distance calculations were performed using the python 3.7 language.

### Odor source search algorithm

The details of the three algorithms for which simulation experiments were performed are described here.

#### surge-zigzagging

A surge-zigzagging algorithm models the female search behavior of an adult male silkmoth, which is a walking insect (Figure 5-Figure supplement 1A). In response to a one-shot sex pheromone stimulus, the silkmoth exhibits a movement that advances in the direction of the odor and a straight movement (surge), followed by a rotational movement (zigzag/loop) to search for further odor information from all directions. The flow chart of the surge-zigzagging algorithm is shown in the supplementary figures. The surge state in the algorithm lasts for 0.5 seconds because surge behavior lasts for about 0.5 seconds after the odor stimulus is presented. Because the surge behavior is elicited when the odor stimulus is presented, the trajectory becomes linear when the odor is continuously (high frequency) presented.

#### casting

A casting algorithm models the odor source localization behavior of a flying moth by mapping it onto a two-dimensional plane (***Li et al., 2001***). A schematic diagram of this algorithm and an implemented flowchart are shown in the supplementary figure (Figure 5-Figure supplement 1B). When the agent in this algorithm detects an odor plume, it moves upwind, and when it loses sight of the plume, it moves in the crosswind direction to rediscover the plume. At this time, when moving upwind, it moves at a certain angle *β* with respect to the upwind direction and continues moving a certain distance *d*_*lost*_ after losing sight of the plume. By repeating these movements in the upwind and crosswind directions, the odor source is localized. The parameters of *β* and *d*_*lost*_ are 30°and 2.5 cm, respectively, and these coincide with the highest search success rates in previous research (***Lochmatter et al., 2008***).

## Acknowledgments

We thank Dr. Takeshi Sakurai (Tokyo University of Agriculture) for providing the sex pheromone, bombykol.

## Additional information

### Funding

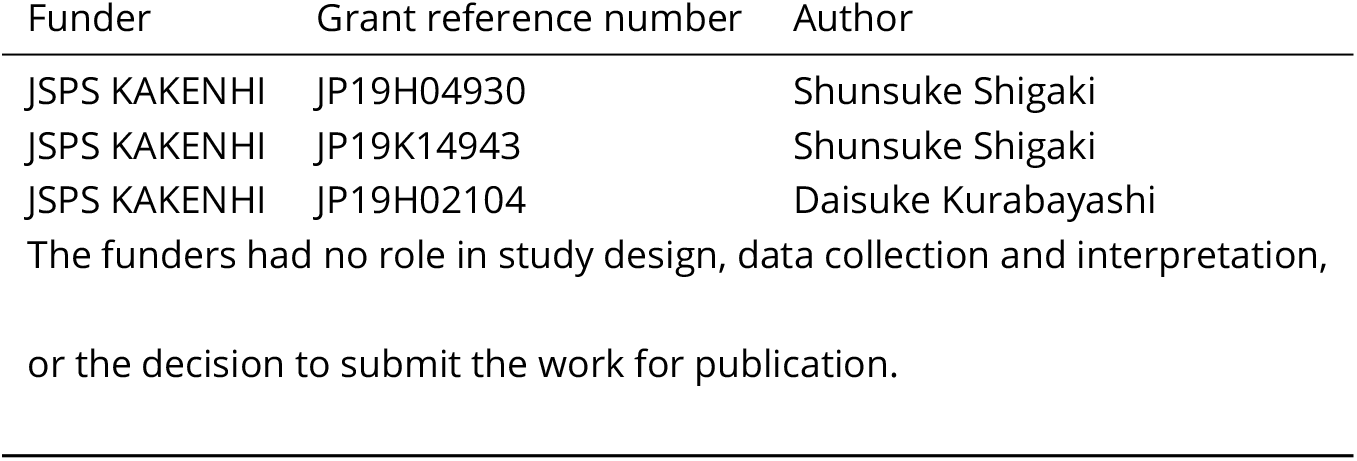

### Author contributions

MY, OH and SS, Planned and designed experiments, Conducted experiments, Analysed and interpreted data, Conception and design, Acquisition of data, Drafting or revising the article; KH and DK, Analysed and interpreted data, Conception and design, Drafting or revising the article

### Ethics

Animal experimentation: Study involved experiments on silkmoths and were conducted according to ethical guidelines

---

**Figure 1–Figure supplement 1.**
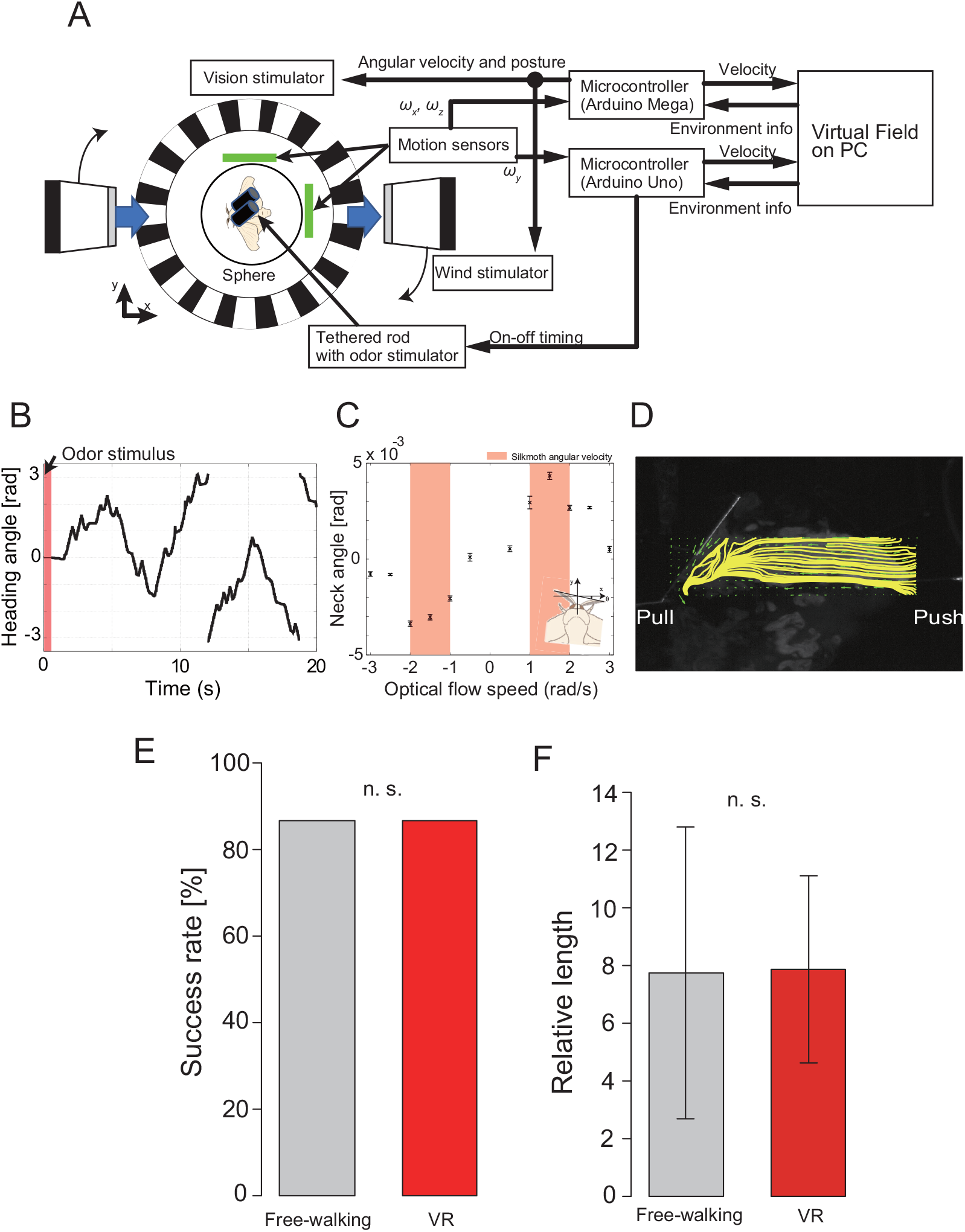
Data comparison between free walking experiment and VR experiment. A: VR system configuration diagram. Two microcomputers were used to control the stimulator and measure the amount of rotation of the sphere. B—D: Evaluation experiment results of each stimulator. B is a graph showing the change in the heading angle of the silkmoth. It was confirmed that when the odor was presented from the upper part of the antennae, the female search behavior was elicited while changing the heading angle significantly. C investigated whether the visual motion reflex was elicited by the visual stimulator by observing the change in the angle of the neck. Because the neck tilt was the largest in the same speed band as the angular velocity of the silk moth, it was confirmed that the visual stimulus could be presented correctly. D is a snapshot of the push-pull rectifier, which is a wind stimulator, visualized by PIV. Since the yellow line represents the streamline and does not generate vortices, it was confirmed that rectification can be generated by push-pull. E: Comparison of search success rates between free walking experiments and VR experiments (Fisher’s exact test, *p <* 0.05). The free walking experiment is the result of two repeated experiments using 15 silkmoths (*n* = 30). The environment of the free walking experiment was set to be the same as the virtual environment of the VR system. F: Comparison of a relative length. The relative length is an evaluation value that standardizes the distance actually traveled by the shortest distance to the odor source. This makes it possible to evaluate how much action was taken in response to sensory input. Because there was no difference between free walking and VR (Welch’s t-test, *p <* 0.05), it is highly possible that the search behavior expressed by a silkmoth in the VR experiment is the same as the behavior in the free walking experiment.

**Figure 5–Figure supplement 1.**
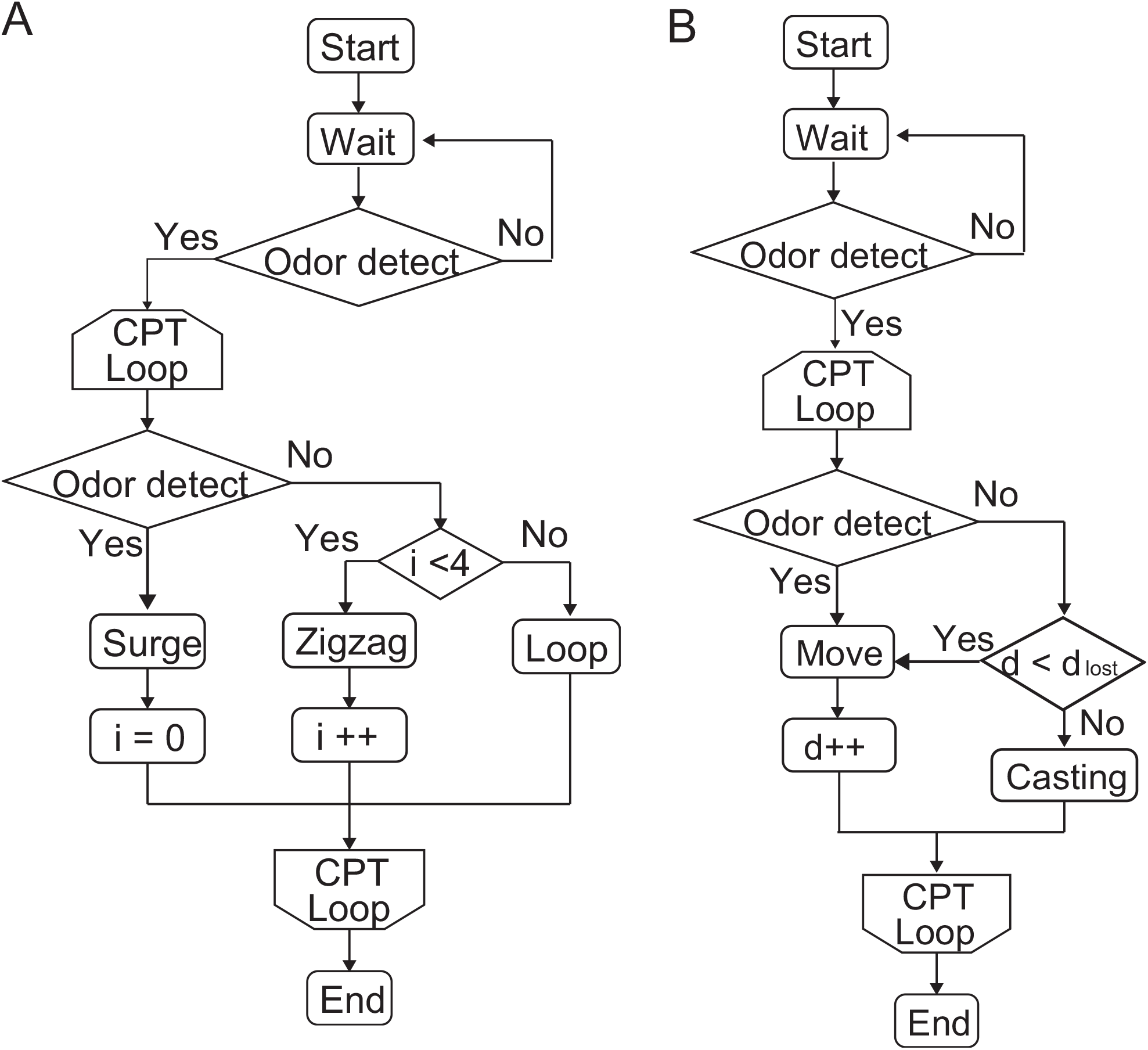
Flowchart of the algorithm used in the simulation experiment. A: Flowchart of the surge-zigzagging algorithm. The surge state and the zigzag/loop state are switched by the odor stimulus. B: Flowchart of the casting algorithm. By detecting the odor, it moves upwind, and when it loses its odor, it casts in the crosswind direction.

## Notes

### Competing Interest Statement

The authors have declared no competing interest.

## References

Adrian RJ. Twenty years of particle image velocimetry. Experiments in fluids. 2005; 39(2):159–169.

Berg HC. E. coli in Motion. Springer Science & Business Media; 2008.

Celani A. Olfactory Navigation: Tempo is the key. Elife. 2020; 9:e63385.

Chen Xx, Huang J. Odor source localization algorithms on mobile robots: A review and future outlook. Robotics and Autonomous Systems. 2019; 112:123–136.

Crimaldi J, Koseff J. High-resolution measurements of the spatial and temporal scalar structure of a turbulent plume. Experiments in Fluids. 2001; 31(1):90–102.

Demir M, Kadakia N, Anderson HD, Clark DA, Emonet T. Walking Drosophila navigate complex plumes using stochastic decisions biased by the timing of odor encounters. Elife. 2020; 9:e57524.

Dyhr JP, Higgins CM. The spatial frequency tuning of optic-flow-dependent behaviors in the bumblebee Bombus impatiens. Journal of Experimental Biology. 2010; 213(10):1643–1650.

Dyhr JP, Morgansen KA, Daniel TL, Cowan NJ. Flexible strategies for flight control: an active role for the abdomen. Journal of Experimental Biology. 2013; 216(9):1523–1536.

Fujiwara T, Kazawa T, Sakurai T, Fukushima R, Uchino K, Yamagata T, Namiki S, Haupt SS, Kanzaki R. Odorant concentration differentiator for intermittent olfactory signals. Journal of Neuroscience. 2014; 34(50):16581–16593.

González E, Marzal F, Miñana A, Doval M. Influence of exhaust hood geometry on the capture effciency of lateral exhaust and push–pull ventilation systems in surface treatment tanks. Environmental progress. 2008; 27(3):405–411.

Haberkern H, Hedwig B. Behavioural integration of auditory and antennal stimulation during phonotaxis in the field cricket Gryllus bimaculatus. Journal of Experimental Biology. 2016; 219(22):3575–3586.

Hernandez-Reyes CA, Fukushima S, Shigaki S, Kurabayashi D, Sakurai T, Kanzaki R, Sezutsu H. Identification of exploration and exploitation balance in the silkmoth olfactory search behavior by information-theoretic modeling. Frontiers in Computational Neuroscience. 2021; 15.

Kanzaki R. Coordination of wing motion and walking suggests common control of zigzag motor program in a male silkworm moth. Journal of Comparative Physiology A. 1998; 182(3):267–276.

Kaushik PK, Renz M, Olsson SB. Characterizing long-range search behavior in Diptera using complex 3D virtual environments. Proceedings of the National Academy of Sciences. 2020; 117(22):12201–12207.

Kikas T, Ishida H, Webster DR, Janata J. Chemical plume tracking. 1. Chemical information encoding. Analytical chemistry. 2001; 73(15):3662–3668.

Li W, Farrell JA, Card RT. Tracking of fluid-advected odor plumes: strategies inspired by insect orientation to pheromone. Adaptive Behavior. 2001; 9(3-4):143–170.

Lochmatter T, Raemy X, Matthey L, Indra S, Martinoli A. A comparison of casting and spiraling algorithms for odor source localization in laminar flow. In: 2008 IEEE International Conference on Robotics and Automation IEEE; 2008. p. 1138–1143.

Lockery SR. The computational worm: spatial orientation and its neuronal basis in C. elegans. Current opinion in neurobiology. 2011; 21(5):782–790.

Minegishi R, Takashima A, Kurabayashi D, Kanzaki R. Construction of a brain–machine hybrid system to evaluate adaptability of an insect. Robotics and Autonomous Systems. 2012; 60(5):692–699.

Murlis J, Elkinton JS, Carde RT. Odor plumes and how insects use them. Annual review of entomology. 1992; 37(1):505–532.

Murlis J, Willis MA, Cardé RT. Spatial and temporal structures of pheromone plumes in fields and forests. Physiological entomology. 2000; 25(3):211–222.

Naik H, Bastien R, Navab N, Couzin ID. Animals in Virtual Environments. IEEE Transactions on Visualization and Computer Graphics. 2020; 26(5):2073–2083. doi: 10.1109/TVCG.2020.2973063.

Obara Y. Bombyx mori mationg dance: An essential in locationg the female. Applied Entomology and Zoology. 1979; 14(1):130–132.

Pansopha P, Ando N, Kanzaki R. Dynamic use of optic flow during pheromone tracking by the male silkmoth, Bombyx mori. Journal of Experimental Biology. 2014; 217(10):1811–1820.

Renou M. Pheromones and general odor perception in insects. Neurobiology of chemical communication. 2014; 1:23–56.

Rubner Y, Guibas LJ, Tomasi C. The earth moverfs distance, multi-dimensional scaling, and color-based image retrieval. In: Proceedings of the ARPA image understanding workshop, vol. 661; 1997. p. 668.

Russell RA, Bab-Hadiashar A, Shepherd RL, Wallace GG. A comparison of reactive robot chemotaxis algorithms. Robotics and Autonomous Systems. 2003; 45(2):83–97.

Ryohei K, Naoko S, Tatsuaki S. Self-generated zigzag turning of Bombyx mori males during pheromonemediated upwind walking. Zoological science. 1992; 9(3):515–527.

Schmitz B, Scharstein H, Wendler G. Phonotaxis in Gryllus campestris l.(orthoptera, gryllidae). Journal of comparative physiology. 1982; 148(4):431–444.

Shigaki S, Fukushima S, Kurabayashi D, Sakurai T, Kanzaki R. A novel method for full locomotion compensation of an untethered walking insect. Bioinspiration & biomimetics. 2016; 12(1):016005.

Shigaki S, Haigo S, Reyes CH, Sakurai T, Kanzaki R, Kurabayashi D, Sezutsu H. Analysis of the role of wind information for effcient chemical plume tracing based on optogenetic silkworm moth behavior. Bioinspiration & biomimetics. 2019; 14(4):046006.

Shigaki S, Sakurai T, Ando N, Kurabayashi D, Kanzaki R. Time-varying moth-inspired algorithm for chemical plume tracing in turbulent environment. IEEE Robotics and Automation Letters. 2017; 3(1):76–83.

Shigaki S, Shiota Y, Kurabayashi D, Kanzaki R. Modeling of the Adaptive Chemical Plume Tracing Algorithm of an Insect Using Fuzzy Inference. IEEE Transactions on Fuzzy Systems. 2019; 28(1):72–84.

Wehner R. Desert ant navigation: how miniature brains solve complex tasks. Journal of Comparative Physiology A. 2003; 189(8):579–588.

Wyatt TD. Pheromones and animal behavior: chemical signals and signatures. Cambridge University Press; 2014.

Yanagawa SS Ryota, Shiota, Kurabayashi D. Construction of chemical plume tracing simulator in a nonrectifying environment. In: IEEE; 2018. p. 147–150.

